# A Single Molecule Investigation of I-Motif: Stability, Folding Kinetics, and Potential as an In-situ pH Sensor

**DOI:** 10.1101/2021.12.17.473157

**Authors:** Golam Mustafa, Prabesh Gyawali, Jacob A. Taylor, Parastoo Maleki, Marlon V. Nunez, Michael C. Guntrum, Hamza Balci

**Affiliations:** Department of Physics, Kent State University, Kent, OH 44242

## Abstract

We present a collection of single molecule work on the i-motif structure formed by the human telomeric sequence. Even though it was largely ignored in earlier years of its discovery due to its modest stability and requirement for physiologically low pH levels (pH<6.5), the i-motif has been attracting more attention recently as both a physiologically relevant structure and as a potent pH sensor. In this manuscript, we establish single molecule Förster resonance energy transfer (smFRET) as a tool to study the i-motif over a broad pH and ionic conditions. We demonstrate pH and salt dependence of i-motif formation under steady state conditions and illustrate the kinetics of i-motif folding in real time at the single molecule level. We also show the prominence of intermediate folding states and reversible folding/unfolding transitions. We present an example of using the i-motif as an in-situ pH sensor and use this sensor establish the time scale for the pH drop in a commonly used oxygen scavenging system.

## Introduction

Depending on the nucleic acid sequences and environmental conditions, DNA can form secondary structures other than the canonical double helix structure. These non-canonical structures include G-quadruplex (GQ) and intercalated motif (i-motif) which form depending on temperature, type and concentration of cations present, and pH ^1–8^. The cytosine rich (C-rich) sequences can adopt i-motif structure that is formed by two parallel duplexes which are held together by intercalated cytosine-hemiprotonated cytosine (C:CH^+^) base pairs ^1,9–11^. The structure also has three loop regions that are unstructured (Figure 1A).

**Figure 1.**
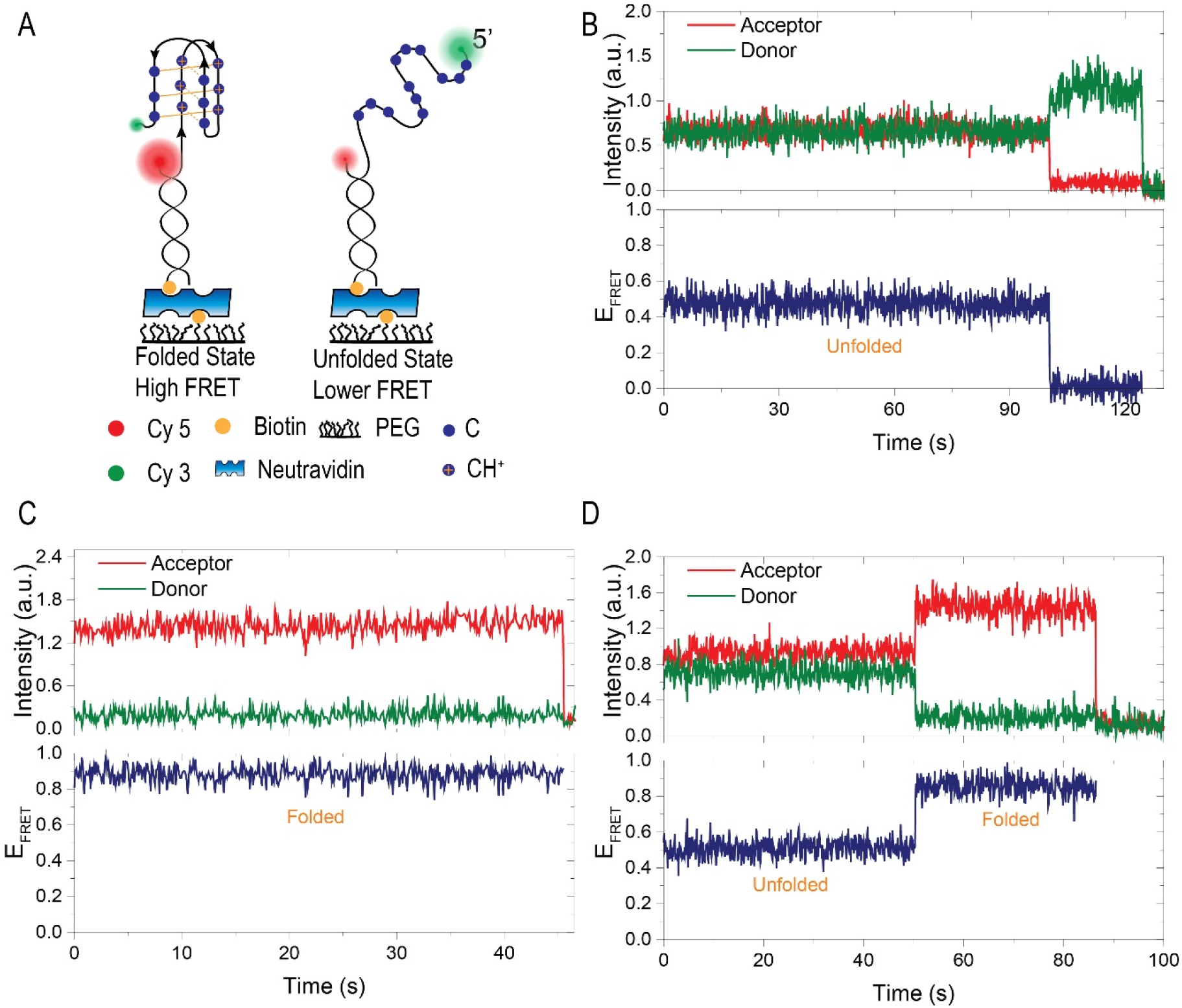
A schematic of the DNA construct and an example smFRET time trace. (A) A partial duplex DNA construct that has an acceptor (Cy5) on the short strand and a donor (Cy3) at the free-end of the long strand. Folding of the DNA into i-motif brings the donor-acceptor fluorophores closer to each other and results in an increase in the FRET efficiency. (B)-(D): Example smFRET traces that show unfolded state (B), folded state (C), and a transition from the unfolded state to folded state (D).

Biophysical studies revealed that the stability of the i-motif depends on the sequence^7^ and length of the loops where i-motif with shorter loops are more stable ^12^. The stability also depends on the environmental condition such as pH ^13^, temperature ^8^, ionic strength ^8,14,15^, cationic effects ^15–17^ and molecular crowding ^18,19^. The structure of i-motif DNA in solution at various pH conditions was investigated by using synchrotron small-angle X-ray scattering technique ^13^.

As protonation is required for C:CH^+^ base pair formation, the i-motif structure shows a sharp transition from folded i-motif to random coil (unfolded) conformation around pH 6.5, below which it remains folded ^9–11,20,21^. Formation, stability, and kinetics of i-motif structures at physiological pH are not only of fundamental interest, but also potentially significant for sensor applications. Several studies have now established that it is possible to stabilize i-motif structures at physiological pH under molecular crowding conditions ^18,22,23^, in a negative supercoiling DNA template ^24^, in the presence of silver(I) cations ^25^, or in the presence of carboxyl-modified single walled carbon nanotubes ^26^. The reversible and fast transformability of the C-rich sequences with pH, from unfolded to a folded i-motif, has potential applications in designing DNA nanomachines ^10,27,28^, logic operation switches ^29–31^, electrochemical sensors for proton detection ^32^ and cellular pH sensors ^33,34^. By introducing structural changes within the DNA constructs, it is possible to tune the pH range that can be sensed by the i-motif and make it a more versatile pH sensor.

Moreover, the i-motif structure has been observed in the nuclei of human cells ^35^, which gives significant support for the physiological relevance of these structures. C-rich sequences that can fold into i-motif are found in telomeres ^36^, in centromeres ^37^, and near the promoter region of oncogenes ^38–40^. Formation of i-motif has been shown to influence DNA replication ^41^ and transcription ^42–45^. For an extensive description of the roles of i-motif structures in the cell, we refer the reader to an excellent recent review^46^.

Folding and stability of the i-motif have been studied as a function of pH and ionic conditions using bulk biophysical methods^8,9,46^. However, limited work has been performed using single molecule methods^47–50^. In this study, we used single molecule Förster resonance energy transfer (smFRET) to investigate stability, kinetics, intermediate folding states, and evolution of i-motif formation at different pH and ionic conditions. After introducing the construct and the method in Figure 1, we demonstrate the capabilities of our approach in Figure 2 and Figure 3 clearly identifying the i-motif structure and evolution of its folding over a broad pH and salt concentration range. In Figure 4, we took advantage of our single molecule approach to monitor the folding dynamics of i-motif in real time using pH jump experiments. These measurements clearly demonstrate the presence of at least one prominent intermediate folding state that is populated during most of folding events. In Figure 5 and Figure 6, we present proof-of-principle measurements that demonstrate the capability of the i-motif structure as an in-situ pH sensor for smFRET studies. We demonstrate that the i-motif structure can sense the time-dependent drop in pH of the environment when “gloxy” (glucose oxidase plus catalase), the most commonly used oxygen scavenging system for single molecule fluorescence studies, is used to enhance photostability of the dyes.

**Figure 2.**
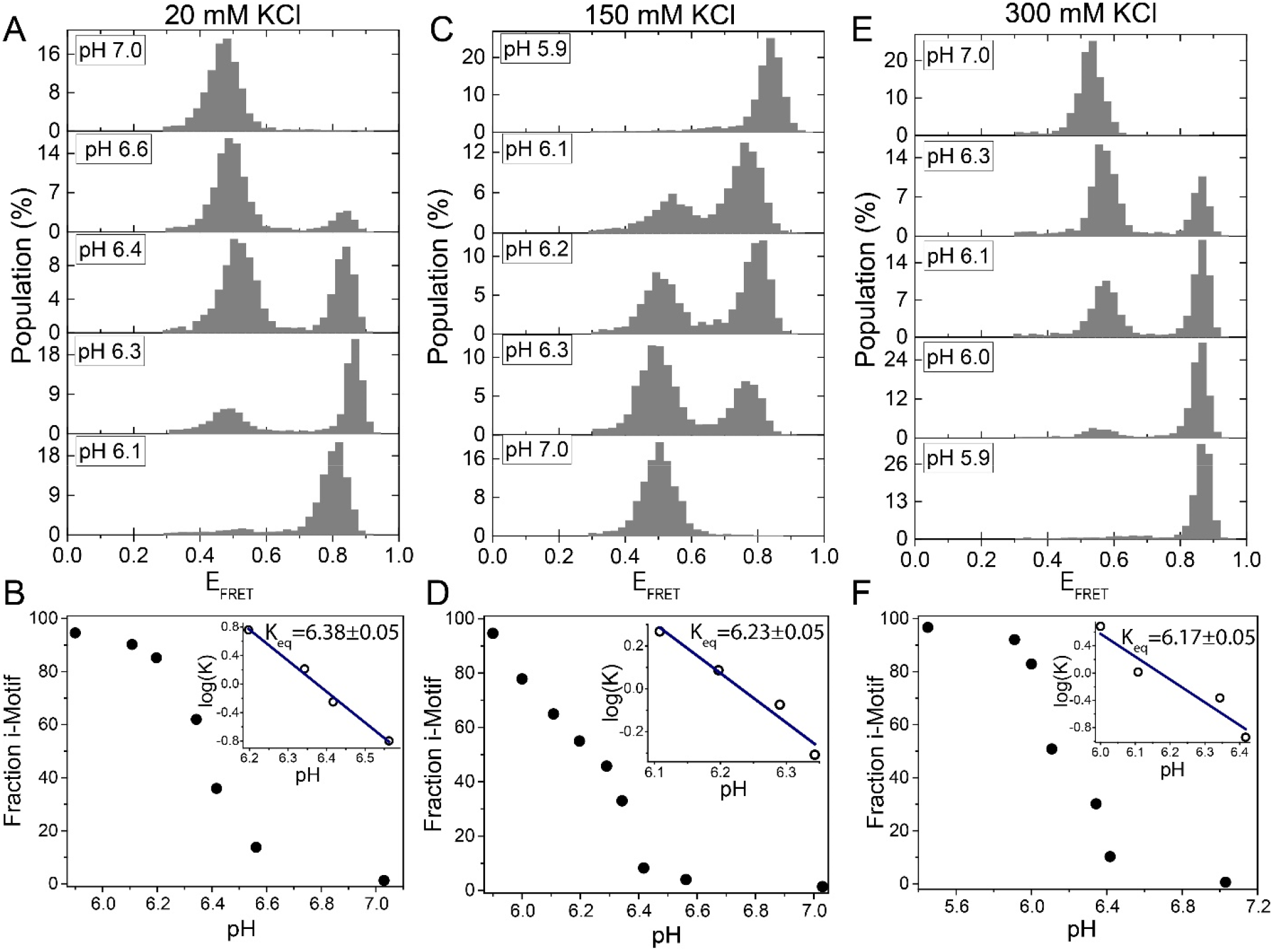
Titration of pH at a constant KCl concentration. The pH was titrated from 7.0 to 5.9 for (A)-(B) 20 mM KCl; (C)-(D) 150 mM KCl; (E)-(F) 300 mM KCl. Under all these ionic conditions, the DNA molecules are exclusively unfolded at pH 7.0 while they are exclusively folded at pH 5.9. At intermediate pH values around pH 6.3, we observe two peaks that represent the folded or unfolded states. (B), (D), and (F) show the evolution of the fraction of i-motif molecules as a function of pH. Blue lines in the insets are linear fits to the most rapidly varying segments of the curve of log(K) vs. pH data.

**Figure 3.**
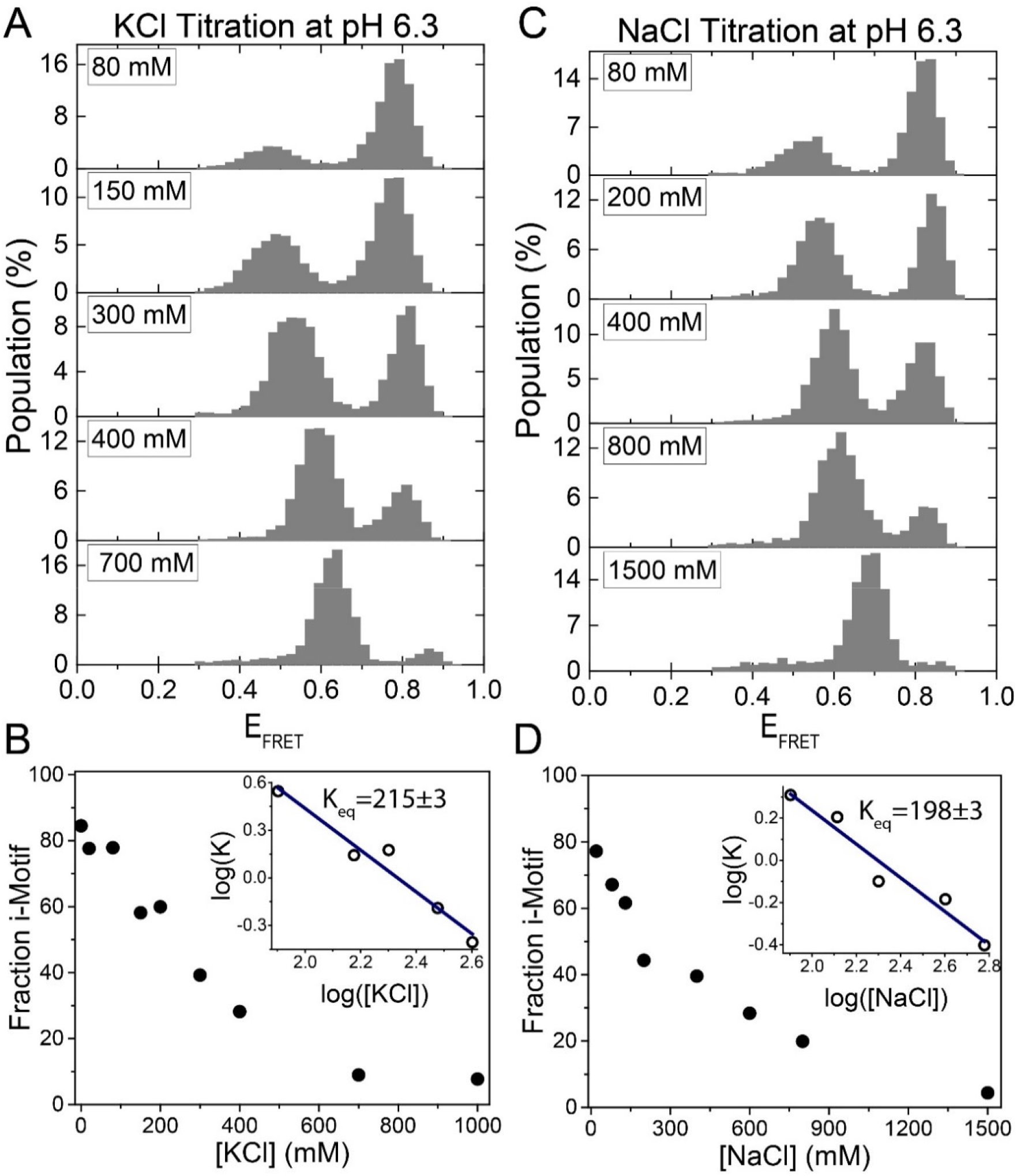
Titration of KCl or NaCl at pH 6.3. KCl was titrated between 0-1000 mM shown in (A) and (B) while NaCl titration is shown in (C) and (D). While most molecules are folded at low salt concentration, they gradually unfold as the salt concentration is increased. (B) and (D) show fraction of i-motif molecules as a function of [KCl] or [NaCl], respectively. Blue lines in the insets are linear fits to the most rapidly varying segments of the curve of log(K) vs. log(salt concentration) data.

**Figure 4.**
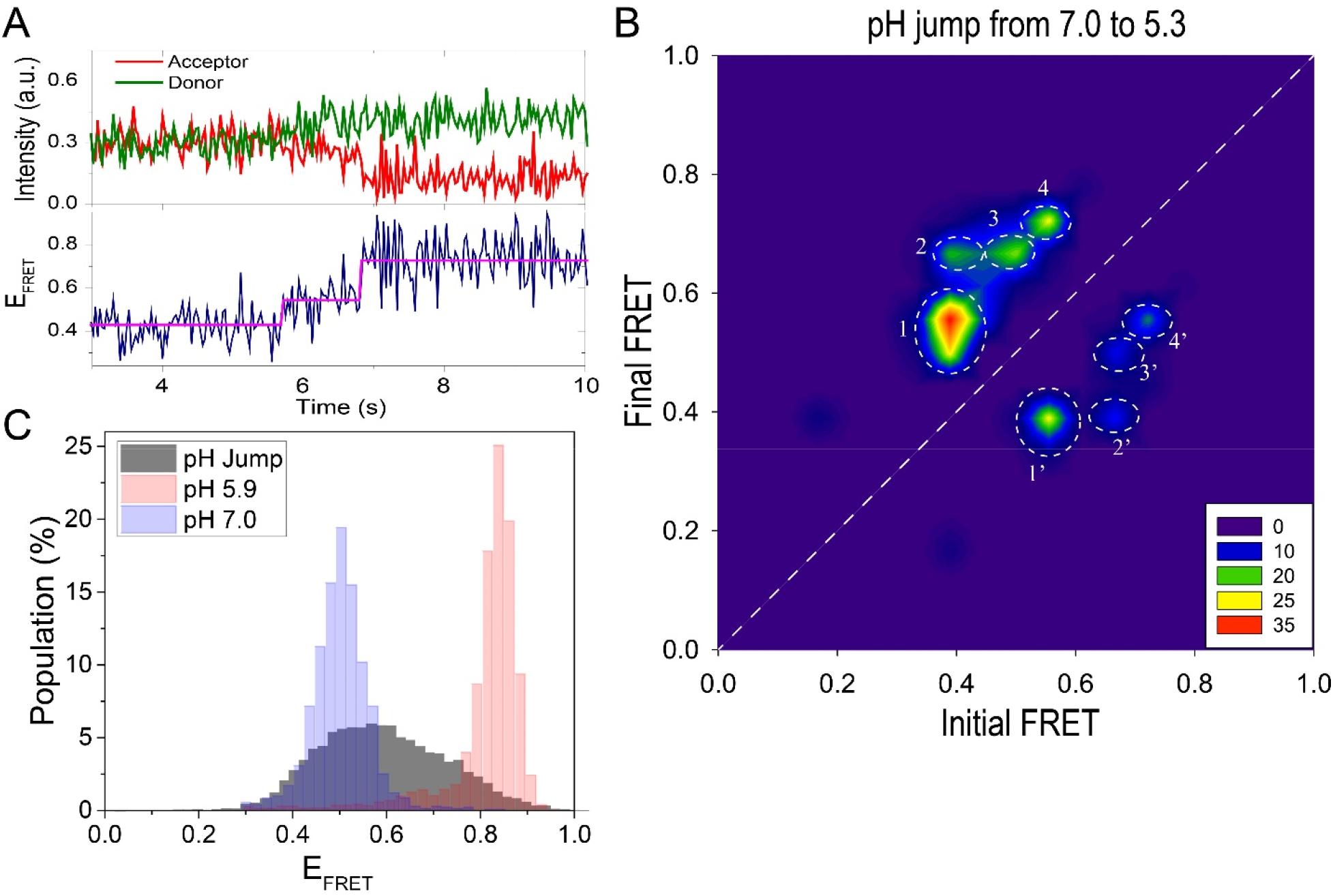
Real time monitoring of the folding process using pH jump measurements. (A) An example time trace is shown. The DNA molecules transition from an unfolded to a folded state as the pH is lowered from 7.0 to 5.3. The trace shows a clear intermediate state that was detected with HMM analysis (the magenta line overlaid on the data). (B) The TDP shows a contour plot of transitions from 104 molecules. The prominent transitions are marked with white circles and numbered as 1,2,3, and 4 in the folding direction. The reverse of these transitions in the unfolding direction are numbered as 1’, 2’, 3’, and 4’. All folding and unfolding processes go through intermediate states between the completely unfolded E_FRET_ ≈ 0.40 and completely folded E_FRET_ ≈ 0.75 states. (C) A histogram (gray) constructed from the FRET levels spanning the interval 2 s before and 2 s after the transition. This representation amplifies the contribution of intermediate states which are not present in steady state histograms obtained 30-min after the buffer exchange is made. For reference, the steady state distributions at pH 7.0 (completely unfolded) and pH 5.9 (completely folded) are shown with blue and red histograms, respectively.

**Figure 5.**
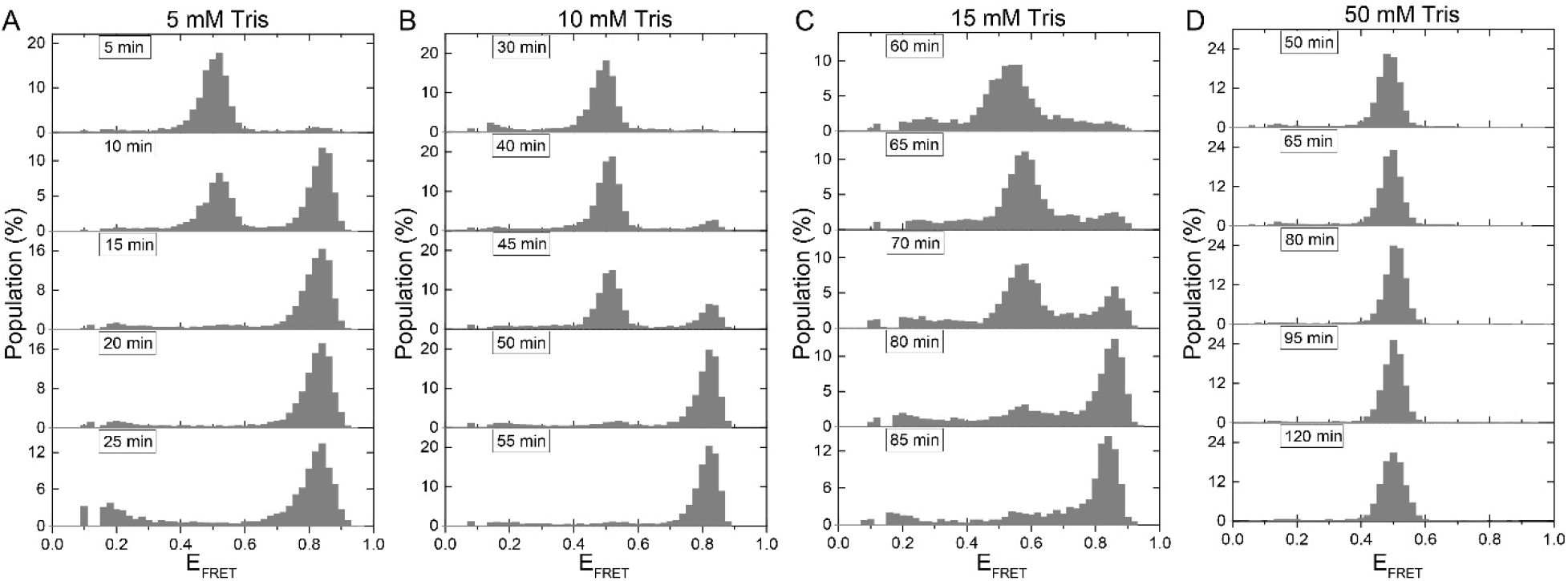
The use of i-motif as an in-situ pH sensor for smFRET experiments. The gradual acidification of the environment when gloxy is used as the oxygen scavenging system is quantified for: (A) 5 mM Tris; (B) 10 mM Tris; (C) 15 mM Tris; and (D) 50 mM Tris. As the buffer strength is increased, the time it takes for the pH to drop from 7.5 to about 6.5, where i-motif folding starts, increases. At 50 mM Tris, all the DNA molecules remain unfolded throughout the 120-minute imaging period.

**Figure 6.**
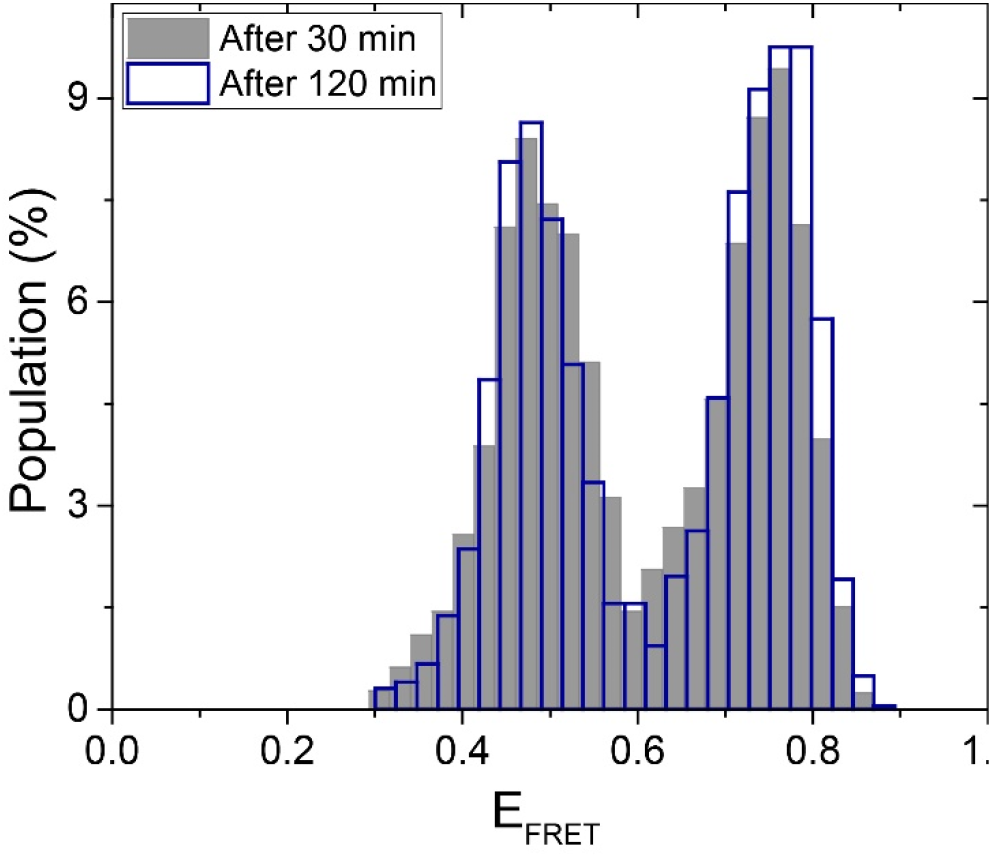
The stability of pH in PCA/PCD oxygen scavenging system. The FRET distribution essentially remains unchanged over 120-minute of imaging when PCA/PCD is used as the oxygen scavenging system, suggesting the pH remains constant to within ±0.05 of the initial pH of 6.3. The gray and blue histogram represent the distributions 30 min and 120 min, respectively, after incubation of PCA/PCD in the chamber.

## Materials and Methods

### DNA Constructs

The following modified (fluorescently labeled and biotin conjugated) DNA oligonucleotides were purchased as HPLC purified from Integrated DNA Technologies (Coralville, IA, USA): Stem Strand: / Biotin / GCCTCGCTGCCGTCGCCA-Cy5 i-motif Strand: Cy3-TT<u>CCCTAACCCTAACCCTAACCC</u>TTTGGCGACGGCAGCGAGGC The underlined nucleotides form the i-motif structure. A partial duplex DNA (pdDNA) construct that has an 18-bp duplex stem and a single-stranded overhang, which forms the i-motif, was prepared by annealing the stem strand and the i-motif strand at 90 °C for 3 min followed by cooling to room temperature over 3 hours. The annealing was performed in 150 mM KCl and 10 mM 2-ethanesulfonic acid, also known as MES, at pH 7.0.

### Sample preparation and smFRET assay

Laser-drilled quartz slides and glass coverslips were cleaned, including a piranha etching step, and coated with polyethylene-glycol (PEG) to prevent non-specific binding of DNA molecules to the surface. A mixture containing a ratio of 100:2 of m-PEG-5000: biotin-PEG-5000 (purchased from Laysan Bio Inc.) was used to coat the surface. The sample chamber was then prepared by sandwiching a double-sided tape between these PEGylated slides and coverslip and treated with 0.01mg/ml neutravidin, which enabled immobilization of biotin-tagged pdDNA constructs on the biotin-PEG surface. This surface cleaning and DNA immobilization steps are similar to those reported previously ^51^.

A solution of 50 pM pdDNA was diluted from the 1 μM annealing stock in multiple steps and was incubated in the sample chamber for 3-5 min at 150 mM KCl and 10 mM MES (pH 7.0). The chamber was then washed with 10 mM MES (pH 7.0) to remove excess pdDNA. Approximately 250-280 molecules/imaging area (~4 × 10^3^ μm^2^) were obtained with this procedure. For all measurements other than the pH jump experiments presented in Figure 4, 20 short movies of 15 frames/movie were recorded at frame acquisition rate of 100 ms/frame for each assay condition. Longer movies (2000-4000 frames at 31 ms/frame) were recorded for pH jump experiments where folding dynamics was studied.

We used two different imaging solutions for our measurements. For all the data, except those presented in Figure 4, the imaging solution contained 50 mM MES of indicated pH, 2 mM Trolox, 25 mM protocatechuic acid (PCA), 0.35 mg/ml protocatechuate-3,4-dioxygenase (PCD), 0.1 mg/ml bovine serum albumin (BSA), 2 mM MgCl_2_, and indicated concentrations of KCl. This solution will be referred to as PCA/PCD imaging buffer. When preparing the buffers, we ensured the pH is stable to three decimal points, but for conciseness of the reporting, we rounded these to the nearest one decimal point, e.g. pH 6.324 was reported as 6.3. The only exception to this is when equilibrium constants were reported. For these, we reported two decimal points, e.g. 6.32, to demonstrate the variation between different conditions within the uncertainties of the measurement (±0.05 or less in pH). Trolox increases brightness of the fluorophores by quenching their dark triplet state. PCA and PCD form an oxygen scavenging system that delays photobleaching of fluorophores. BSA patches the surface areas that might have imperfect PEGylation. To allow the system to reach steady state and PCA/PCD to reduce the oxygen concentration to adequate levels, the imaging solution was incubated with the DNA molecules for 30 min before measurement.

For the pH jump measurements presented in Figure 4 glucose oxidase and catalase (also called gloxy) was used as the oxygen scavenging system instead of the PCA/PCD system. The contents of the imaging buffer were otherwise same as listed above. This solution will be referred to as gloxy imaging buffer. A significant advantage of gloxy, compared to PCA/PCD, is that long incubation times are not required to remove oxygen, and data acquisition can effectively start as soon as the imaging buffer is introduced into the chamber. This enables observing folding of the i-motif in real time as movies could be recorded while pH of the environment was dropped from 7.0 to 5.3 via a buffer exchange (called pH jump in this study and “flow measurement” in some others where structural changes in DNA were recorded in real time^52^). For these measurements, the DNA molecules were first incubated in a PCA/PCD imaging buffer that contains 50 mM MES at pH 7.0 and 150 mM KCl. After 30 min of incubation, recording of a long movie of 2000-4000 frames at an acquisition rate of 31 ms/frame was initiated. After recording ~200 frames, a gloxy imaging buffer containing 50 mM MES at pH 5.3 (all other ingredients were kept identical) was flown into the sample chamber, while the recording continued.

Even though gloxy made the pH jump experiments possible, it is known to result in a gradual pH drop in the chamber, due to generation of gluconic acid as the by-product of reducing the free oxygen concentration according to the following reaction:

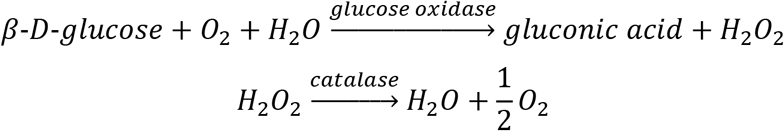

The rate of the pH drop depends on the buffer strength, which we quantified in Figure 5 by using the i-motif as an in-situ pH sensor. These measurements started from physiological pH (7.5) which motivated the use of Tris-HCl as the buffer instead of MES due to its higher pKa^53^. The imaging solution contained the following for these measurements: Indicated concentration of Tris-HCl (initial pH 7.5), 0.1 mg/ml glucose oxidase, 0.02 mg/ml catalase, 2 mM Trolox, 0.1 mg/ml BSA, 2 mM MgCl_2_, and 150 mM KCl.

### Imaging Setup

A prism-type total internal reflection fluorescence microscope equipped with an Olympus IX-71 microscope and an Andor IXON EMCCD camera (IXON DV-887 EMCCD, Andor Technology, CT, USA-now part of Oxford Instruments) was used for the measurements. The donor fluorophore was excited with 532 nm wavelength green laser (Spectra Physics Excelsior). An Olympus water objective (60x, 1.20 NA) was used for collecting the fluorescence signal.

### Data Analysis

Using a custom software written in C++, the recorded movies were analyzed and time traces of donor intensity (I_D_) and acceptor intensity (I_A_) for each molecule were generated. The FRET efficiency (E_FRET_) was calculated using *E_FRET_* = *I_A_*/(*I_A_* + *I_D_*. The FRET efficiency population histograms were constructed from single molecule traces such that each molecule contributed equally, regardless of how long the molecules remained fluorescent (time it took for molecules to photobleach). The leakage of the donor signal into the acceptor channel, which results in the donor-only (DO) peak, was subtracted from the histograms and the DO E_FRET_ level was shifted to zero to correct for this leakage.

In order to analyze the pH and salt concentration dependence of the i-motif structure, we utilized a published method^7^. Accordingly, an equilibrium constant *K* is defined as:

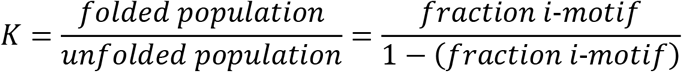

When the folded and unfolded populations are equal *K* = 1. The pH or salt concentration where *K* = 1 is satisfied will be referred to as *K_eq_*. To determine *K_eq_*, we plot *log*(*K*) *vs*. pH or *log*(*K*) *vs*. *log*(*salt concentration*), and linearly fit the most rapidly varying segments of the curve. When *K* = 1 log(*K*) = 0 so pH or salt concentration where the line crosses 0 will be the corresponding K_eq_. The slope of *log*(*K*) *vs*. pH has been interpreted as the number of protons gained or lost during the folding or unfolding transitions, respectively ^7^.

Origin software program was used for these analyses. The errors in the cited *K_eq_* values are the standard deviations obtained from the fitting analysis.

## Results and Discussion

### I-motif stability as a function of pH

In this section, we present smFRET studies that demonstrate the capabilities of the method to distinguish between the folded and unfolded i-motif structures under different pH and ionic conditions. Figure 1A shows a schematic of the DNA construct and the donor-acceptor dye positions. Figure 1B-D show example smFRET time traces that show unfolded state at FRET efficiency E_FRET_ = 0.5 (Figure 1B), folded state at E_FRET_ =0.9 (Figure 1C) and a transition from the unfolded to the folded state where the FRET efficiency rises from E_FRET_ ≈ 0.5 to E_FRET_ ≈ 0.9 (Figure 1D). Histograms of time traces, each containing data from several thousand molecules, show two well-separated peaks representing the folded and unfolded states (Figure 2).

Figure 2 shows pH titration (pH 5.9 to 7.0) at 20 mM (Figure 2A-B), 150 mM (Figure 2CD), and 300 mM KCl (Figure 2E-F). Five smFRET histograms are shown for each KCl concentration ([KCl]). In these histograms, the high FRET states (E_FRET_>0.70) represent the folded i-motif and low FRET states (E_FRET_<0.60) represent the unfolded state. As pH is lowered the folded state population increases while the unfolded state population decreases, as expected. For all tested salt conditions, all molecules were folded at pH 5.9 (a single peak at E_FRET_≈0.80) while they were all unfolded at pH 7.0 (a single peak at E_FRET_≈0.50). Between pH 6.0 to 6.6, the entire population transitions from i-motif to random coil, in agreement with earlier reports^9^. The steady state variation of folded state as a function of pH is illustrated in Figure 2B for 20 mM KCl, in Figure 2D for 150 mM KCl, and in Figure 2F for 300 mM KCl. The pH equilibrium constants (K_eq_), where 50% of molecules are folded, were determined by linear fits (blue lines in the inset of cited figures) to *log*(*K*) *vs*. pH data. These fits resulted in the following: K_eq_=6.38±0.05, K_eq_=6.23±0.05 and K_eq_=6.17±0.05 for 20 mM, 150 mM, and 300 mM KCl, respectively. Higher concentrations of K^+^ require lower pH values for i-motif formation, as expected by Le Chatelier’s principle ^8^. The uncertainties were based on the pH range that results in ±5% variation in the fraction of i-motif population. The equilibrium constants we observe (for example K_eq_=6.23±0.05 at 150 mM KCl) are slightly higher than those observed in bulk measurements. For example, circular dichroism measurements reported K_eq_ ≈5.75 and K_eq_ ≈5.85 at 165 mM and 115 mM KCl, respectively, which can be interpolated to K_eq_ ≈5.8 at 150 mM KCl ^8^. The higher K_eq_ in our measurements, where DNA molecules are immobilized on the surface, might be due to surface-induced stabilization of the i-motif observed for some surfaces ^54^.

### I-motif stability as a function of ion concentration

We also studied the fraction of folded i-motif structures as a function of [KCl] as shown in Figure 3A-B. The pH was fixed at 6.3 for these measurements while [KCl] was titrated from 0-1000 mM. As expected, the unfolded state population increases as [KCl] is increased. It was possible to drive the system from almost completely folded to almost completely unfolded between 80-400 mM KCl, in agreement with earlier studies^9^. As Figure 3A shows, the peak that represents the unfolded state gradually shifts to higher FRET values as [KCl] is increased, which is expected due to better shielding of the negatively charged ssDNA backbone at these higher salt concentrations. This results in the folded and unfolded peaks to approach each other since the shift in the folded state is significantly less prominent, which is also expected since unstructured ssDNA forms only a small fraction of the overall construct (two spacer nucleotides on either side of the i-motif). This could potentially prevent smFRET to be as sensitive to i-motif formation at very high salt concentrations (several molar); however, this is not the case until 1 M KCl. Several molar salt concentrations would require very low pH values for i-motif formation, where photostability of fluorophores commonly used in smFRET measurements is typically poor in our experience. Therefore, it would be challenging to study i-motif structure and kinetics using smFRET at such high salt concentrations. Figure 3B shows the evolution of fraction of i-motif population as function of KCl. A linear fit to *log*(*K*) *vs*. *log*([*KCl*]) data, shown in the inset of Figure 3B, resulted in K_eq_=215±3 mM KCl at pH 6.3. Since the i-motif is closely related to GQ, and the GQ stability shows a strong dependence on the ion type^55^, we performed similar salt titration measurements using NaCl (Figure 3C-D). Unlike the GQ, our analysis demonstrated very similar i-motif stabilities for K^+^ or Na^+^, with K_eq_=198±3 for NaCl.

### Monitoring i-motif folding in real time

In order to observe the folding of the i-motif in real time, we performed buffer exchange measurements where a lower pH buffer (pH 5.3) was injected into the sample chamber that contained unfolded molecules at a higher pH (pH 7.0). Gloxy, rather than PCA/PCD, which was used for the data in Figure 2 and Figure 3, was used in these measurements as the oxygen scavenging system in order to image the folding process in real time, which happens within a few seconds of the buffer exchange. The lower pH buffer was injected into the chamber with a syringe pump that minimized the disturbance on the system and allowed continuous imaging of the folding process. Figure 4A shows a representative smFRET time trace that captures the folding process. The trace shows an intermediate folding state, which was detected by Hidden-Markov modeling (magenta line that are overlaid on the data), performed using the vbFRET program^56^. Figure 4B shows the transition density plot (TDP) constructed from 105 such traces. In TDP, the x-axis represents the FRET state before the transition, initial FRET, while the y-axis represents the FRET state after the transition, final FRET. The color in TDP represents the number of transitions, red being the most frequent and blue the least. For example, a folding transition from E_FRET_ ≈ 0.40 to E_FRET_ ≈ 0.70, it will contribute as one unit in TDP at point (0.40, 0.70). Similarly, an unfolding transition from E_FRET_ ≈ 0.70 to E_FRET_ ≈ 0.40 will contribute to the contour plot at point (0.70, 0.40). The white dashed line indicates the 45°. Transitions above this line are due to folding while transitions below it are due to unfolding. As expected, the levels above the 45°-line are more populated as a higher pH buffer was exchanged with a lower pH buffer and the transitions were biased in the folding direction. However, it should be noted that there are clearly detectable states below the 45°-line indicating that the folding process is not irreversible, and some molecules occasionally unfold during this transition period.

Another interesting revelation of the TDP is the presence of prominent intermediate states, which was not observed in steady state histograms. These distinct transitions are marked with white dashed circles, which are numbered as 1,2,3, and 4 in the folding direction. The reverse of these transitions in the unfolding direction are marked with 1’, 2’, 3’, and 4’, respectively. The most populated transitions in TDP are from E_FRET_ ≈ 0.40 to E_FRET_ ≈ 0.55 (circle 1). The molecules that reach E_FRET_ ≈ 0.55 either continue to fold and reach E_FRET_ ≈ 0.70 (Circle 3) or E_FRET_ ≈ 0.75 (Circle 4). There also molecules that directly transition from the unfolded state E_FRET_ ≈ 0.40 to E_FRET_ ≈ 0.70 (Circle 2). All of these transitions are reversible, in the form of transitions shown with circles 1’, 2’, 3’, and 4’. Assuming the E_FRET_ ≈ 0.75 state to be the completely folded state and the E_FRET_ ≈ 0.40 state the completely unfolded state, we do not observe any direct folding or unfolding events, i.e. we do not have transitions around (0.40, 0.75) or (0.75, 0.40). Instead, during both folding and unfolding, the molecules transition to the intermediate state first before they completely fold or unfold. As the intermediate state is a relatively short-lived state that is observed only during the transitions, it is not surprising that it was not detected in steady state histograms depicted in Figures 2 and 3, which survey the molecules after they are incubated in respective buffers for about 30 minutes. In order to amplify the contribution of the intermediate populations in the histograms, we constructed a new histogram from the time traces of these pH jump measurements and included only the segments spanning 2 s before and 2 s after the transition. Figure 4C shows the new histogram where the states between the unfolded (E_FRET_ ≈ 0.40) and folded (E_FRET_ ≈ 0.75) states are significantly populated (gray histogram). For reference, we overlaid the corresponding steady state histograms at pH 5.9 (red histogram) and at pH 7.0 (blue histogram). The population in the gray histogram between the blue and red peaks represents the intermediate folding state visited during the pH jump experiments.

### I-motif as an in-situ pH sensor

The oxygen scavenging system is a critical component of single molecule fluorescence studies that use small organic fluorophores since free oxygen radicals are the primary cause of photobleaching, a chemical transition of the fluorophores to a non-fluorescent state. PCA/PCD and gloxy (glucose oxidase + catalase) are the two most commonly used oxygen scavenging systems and they have different characteristics that make them better choices for different applications, as illustrated in this study. While PCA/PCD provides a constant pH over multiple hours of imaging, it is typically necessary to incubate PCA/PCD for 10-30 minutes in the sample chamber before it becomes effective. On the other hand, gloxy reduces the free oxygen concentration much more rapidly, a few seconds, but it results in accumulation of gluconic acid which gradually reduces the pH. Because of these characteristics, most measurements in this study were performed with PCA/PCD as having a stable pH was essential to avoid having a compounding factor when salt or pH of the environment was changed.

In these measurements, incubating PCA/PCD in the chamber for 30 minutes was possible as we were primarily interested in the steady state distributions. However, in the pH jump measurements where the folding process was much faster and we were interested in the dynamics of this fast process, gloxy was preferred as the oxygen scavenging system. Even though fluorophore brightness and lifetime were demonstrated to be superior in PCA/PCD compared to gloxy for some of the commonly used fluorophores^57^, gloxy is still probably the most widely used oxygen scavenging system in single molecule fluorescence measurements due to this fast action capability. Therefore, it is important to understand the extent of the pH drop in gloxy as a function of time to establish time limits for justifying the assumption of relatively constant pH in the environment.

This important question was previously investigated by using SNARF-1, a ratiometric dual emission dye^58^. By characterizing the relative populations of the two peaks at different pH values, the pH variation within the sample chamber was determined. In this study, the sample chamber was placed in the path of the light beam within the spectrofluorometer and the pH was determined based on bulk FRET signal. In this study, we monitored the pH variation based on folding of the i-motif as function of time. As the pH drop would depend on the strength of the buffer, we performed the same measurement in 5 mM, 10 mM, 15 mM and 50 mM Tris that had an initial pH of 7.5. We would expect the pH drop to be faster in weaker buffers therefore we would expect the initially unfolded DNA strands to fold into i-motif after shorter incubation periods compared to that in stronger buffers. Based on the ionic conditions of this assay (150 mM KCl) and the calibration data presented in Figure 2C-D, we conclude that the pH of the environment should be 5.9 for all the DNA molecules to fold into i-motif.

Figure 5 demonstrates these data where we periodically surveyed different regions of the chamber and quantified the fraction of molecules that remain unfolded or fold into i-motif after introducing gloxy into the chamber. These measurements show that all DNA molecules fold into i-motif after approximately 15 min in 5 mM Tris, 55 min in 10 mM Tris, and 85 min in 15 mM Tris. These data set the time scale for pH to drop from 7.5 to around 6.5, where i-motif population becomes detectable, under these buffer conditions. The DNA molecules remain completely unfolded over 120 minutes of imaging in 50 mM Tris. This suggests that the pH remains above 6.5 (based on Figure 2C-D) in 50 mM Tris even after 120 minutes of incubating the gloxy in the chamber. These results are in quantitative agreement with the pH drop characterized by the ratiometric bulk FRET measurements on SNARF-1^58^. It should be noted that the drop in the pH could be slowed down by using lower concentrations of glucose oxidase; however, this is often accompanied by lower photostability. In our experience, reducing the glucose oxidase concentration by a factor of 10 results in two-fold reduction in Cy3/Cy5 photostability, while fluorophore brightness remains similar. For comparison, we also measured the stability of pH in PCA/PCD over a 120-min imaging period in 150 mM KCl. Since we expect pH to be relatively stable in PCA/PCD, we selected a pH value where even small variations in pH would result in detectable changes in the fraction of the folded i-motif population. Based on the data in Figure 2CD, pH 6.3 would be the ideal choice for this purpose since approximately 50% of the molecules would be folded and small variations in the pH would result in significant changes in the folded fraction. We also used MES buffer for these studies since it is a better buffer in such lower pH studies. Figure 6 shows these data where the histograms after 30-min and 120-min are overlaid. The histograms are essentially identical, suggesting that the pH remains remarkably stable (6.29±0.05) in PCA-PCD over 120-min imaging period.

## Conclusion

Our studies demonstrate that with proper design of the DNA construct and placement of the fluorophores, smFRET can be used to study i-motif systems over a broad range of ionic and pH conditions. Steady state distributions, dynamic transitions, and intermediate states during the folding process are clearly identifiable using different image acquisition approaches. The versatility of the method also suggests that interactions of i-motif structures with proteins, small molecules, or competing nucleic acid sequences would also be possible using smFRET. The i-motif sequence we studied proved to be highly sensitive to pH variations in 6.3±0.3 range. Based on this observation, we demonstrated the feasibility of using the i-motif as an in-situ pH sensor for smFRET experiments to establish the time limits where the pH of the system drops below the physiologically relevant range.

